# Minocycline engages microglia in the midbrain periaqueductal grey to attenuate hypoxia-triggered panic-like behaviour in rats

**DOI:** 10.64898/2026.07.29.741424

**Authors:** Arthur Rocha-Gomes, Jefferson Manoel do Nascimento-Silva, Beatriz Cassimiro Baratella, Beatriz Dominiquini-Moraes, Yurie Sato, Erika Meyer, Sabrina Francesca Lisboa, Norberto Cysne Coimbra, Luciane H. Gargaglioni, Valentina Mosienko, Hélio Zangrossi

## Abstract

Panic disorder (PD) is a chronic and highly disabling psychiatric disorder characterised by recurrent and unexpected panic attacks, with underlying neurobiological mechanisms poorly understood. Emerging evidence suggests that inflammatory processes may contribute to the triggering of panic attacks, with a subset of PD patients exhibiting alterations in circulating cytokine levels, while animal studies indicate that an immunoresponsive microglial phenotype may contribute to the disorder’s pathophysiology. Minocycline, a tetracycline-class antibiotic that crosses the blood-brain barrier, exerts anti-inflammatory effects and has demonstrated therapeutic potential in attenuating panic attacks, supporting its potential as an alternative strategy for reducing PD-related symptoms by modulating microglial activity. In this study, we investigated whether exposure of male Sprague-Dawley rats to a panicogenic stimulus, hypoxia (7% O_2_), is followed by microglial morphological remodelling in the midbrain periaqueductal grey (PAG), a well-known panic-associated structure, at baseline and after minocycline treatment. The effects of hypoxia on the expression of panic-like jumping behaviour were measured during the respiratory challenge, whereas microglial morphology was assessed at 1, 6, or 24h following the aversive stimulus. In a second experiment, the effects of minocycline (30 mg/kg, i.p., administered once daily for 5 days) on the immediate behavioural responses to hypoxia were compared with those produced by an acute administration of alprazolam (2 mg/kg, i.p.), a benzodiazepine widely used in the clinical management of PD. Minocycline effects on microglial morphology in the PAG were also assessed. Our findings show that hypoxia elicited robust jumping behaviour, without affecting overall locomotion. This panicogenic effect was accompanied by marked and time-dependent microglial morphological changes in the dorsomedial (dmPAG) and ventrolateral (vlPAG) columns of the PAG, consistent with a shift towards an immunoresponsive phenotype. Notably, minocycline, similarly to alprazolam, reduced the number of jumps, indicating a panicolytic effect, while also preventing hypoxia-induced microglial remodelling in the dmPAG. Altogether, these findings suggest a role for microglia in regulating hypoxia-induced panic- like behaviour, indicating that microglial inhibition, as achieved here with minocycline, represents a promising therapeutic strategy for preventing panic attacks.

## 1 INTRODUCTION

Panic disorder (PD) is a chronic and disabling psychiatric condition characterized by recurrent and unexpected panic attacks. These attacks consist of abrupt surges of intense fear or discomfort, peaking within minutes and accompanied by at least four of the thirteen physical and/or cognitive symptoms (e.g., palpitations, sweating, trembling, sensations of shortness of breath, chest pain, fear of losing control, or fear of dying) ^1^. Epidemiological data indicate that the lifetime prevalence of PD is approximately 1.7% worldwide ^2^. Notably, an estimated 56.6% of affected individuals experience the disorder with high severity ^3^. Beyond its substantial clinical burden, this condition imposes high economic and societal costs ^4^, highlighting the need to elucidate its etiological mechanisms and to develop novel therapeutic strategies.

An increasing body of evidence also suggests that inflammatory processes may contribute to the pathophysiology of this disorder. Evidence suggests that a subset of patients with PD exhibit elevated systemic levels of pro-inflammatory markers, including interleukin (IL)-1β, IL-6, interferon-γ (IFN-γ), and tumour necrosis factor-α (TNF-α) ^5–7^. Interestingly, our research group recently showed that adult male C57BL/6 mice exposed to hypercapnia (20% CO₂) – a panicogenic condition that evokes a robust escape response in mice ^8^ – exhibited reduced arborisation of brain-resident microglia within the locus coeruleus, a key noradrenergic nucleus central to the regulation of defensive circuits, respiration, antinociception, and autonomic functions ^9,10^. Together, these findings suggest that, in addition to peripheral inflammation, PD may also involve alterations in the microglia immunoresponsive state, supporting a potential role for these cells in the underlying neurobiological mechanisms.

An alternative experimental approach for inducing panic attacks involves exposure to low oxygen concentrations (7% O₂), a paradigm widely used in human studies^11–13^ and previously shown by our group to evoke a robust escape response characterised by upward jumps in rats ^8^. Recent studies have shown that exposure to hypoxia can trigger significant modulation of microglia, leading to inflammatory and neuropathological outcomes in brain regions such as the subfornical organ and cerebral cortex ^14,15^. Microglia are the resident immune surveillance cells of the brain and can detect extracellular pH changes associated with hypoxic exposure. In response to extracellular acidosis, these cells can be rapidly recruited and shift towards an immunoresponsive state aimed at restoring homeostasis ^14,15^. Despite scientific advances in the field, it remains unclear how hypoxia-induced panic-like responses shape the temporal progression of this microglial phenotypic shift and the extent to which these changes disrupt homeostasis in brain regions involved in the expression of panic-related defensive responses.

In the first experiment of this study, we conducted immunofluorescence analyses to assess the temporal effects (1, 6, and 24 hours) of a hypoxic exposure (7% O₂) on microglial morphological changes in male adult rats within the midbrain periaqueductal grey (PAG), a key structure involved in the PD autonomic and behavioural responses ^16,17^. Our findings revealed that hypoxia exposure caused a marked panicogenic-like effect accompanied by microglial morphological changes, with a higher inflammation index in the dorsomedial column of the PAG (dmPAG).

These findings highlight a novel aspect of the hypoxia-induced panic model, in which microglia may not only contribute to neuronal dysfunction but also represent a potential therapeutic target for PD interventions. In this context, we also recently showed that minocycline, a systemically active tetracycline-class antibiotic that crosses the blood-brain barrier ^18–22^, exerts both anti-inflammatory and panicolytic effects in human patients with PD following CO₂ exposure ^9^. Moreover, previous preclinical studies have shown that this compound can reduce glial activation and anxiety-like behaviours, exerting neuroprotective effects when administered preventively ^23^. Therefore, in the second experiment, we evaluated whether minocycline administration inhibits hypoxia-induced microglial alterations within the dmPAG while concurrently reducing the associated panic-like jumping behaviour.

## 2 MATERIAL AND METHODS

### 2.1 Animals and ethics

Male Sprague-Dawley rats were obtained from the animal facility of the University of São Paulo, at Ribeirão Preto Campus (São Paulo, Brazil), and underwent a two-week acclimatisation period prior to the experiment. At the beginning of the experiments, animals were 70-days old and housed in groups of 3-4 in polypropylene cages (39 × 32 × 17 cm) containing autoclaved sawdust under controlled environmental conditions (22 ± 1 °C; 12 h light/dark cycle, lights on at 07:00 h; illumination 60 lux). Standard rodent chow (Nuvilab, Quimtia S/A, Paraná, Brazil) and water were available *ad libitum*. All experimental procedures were conducted in accordance with international guidelines for the ethical use of laboratory animals and were approved by the local Institutional Animal Care and Use Committee (protocol number <u>1134/2022</u>).

### 2.2 Drugs, antibodies and reagents

Minocycline (Hovione Farmaciência, Portugal) and alprazolam (EMS, Brazil) were dissolved in a sterile vehicle, phosphate-buffered saline (PBS; pH 7.4), and prepared freshly before administration. For immunofluorescence assays, primary polyclonal rabbit anti-ionized calcium-binding adapter molecule 1 (Iba1 #019-19741; Fujifilm Wako, Houston, USA) and secondary anti-rabbit Alexa Fluor 594-conjugated (#A-11012; Thermo Fisher Scientific, Houston, USA) antibodies were used.

### 2.3 Apparatus

The apparatus used for the hypoxia challenge consists of a cylindrical Plexiglas arena with transparent acrylic walls (25 x 35 cm), hermetically sealed and equipped with two valves: one at the top, linked to a nitrogen (N_2_) cylinder, and another midway, linked to an air pump. Hypoxia was induced by progressively increasing the N_2_ flow rate to 4.5 L/min, decreasing the oxygen (O_2_) concentration in the chamber to 7%. Chamber O_2_ concentrations were continuously monitored using a gas analyser (ML206 Gas Analyser; AD Instruments, Brazil) and displayed on a PowerLab running Chart 5 software (AD Instruments, Brazil).

### 2.4 Tissue collection and Immunofluorescence

Animals were anaesthetised with ketamine hydrochloride (100 mg/kg) and xylazine (10 mg/kg) and perfused transcardially with PBS (0.1 M; pH 7.3), followed by 4% paraformaldehyde (PFA; Sigma-Aldrich, Brazil). Brains were post-fixed in PFA for 24h, cryoprotected in 30% sucrose for 48h at 4 °C, frozen in dry ice-cooled isopentane at -30 °C, and stored at -80 °C until sectioning. Coronal sections (40 μm) were cut on a cryostat (Leica CM1850, Germany).

For the immunofluorescence analyses, samples were blocked with 5% bovine serum albumin and 0.1% Triton-X100 in PBS for 3h, after which they were incubated with rabbit anti-Iba1 primary antibody (1:500) overnight at 4°C. The following day, sections were triple-washed with 0.1% Triton-X100 in PBS and incubated with Alexa Fluor 594 anti-rabbit IgG secondary antibody (1:1000) for 2h, then mounted using an appropriate mounting medium (Fluormount, Electron Microscopy Sciences, USA).

### 2.5 Image acquisition and microglia morphological analyses

The dorsomedial column of the periaqueductal grey (dmPAG) and the ventrolateral columns of the PAG (vlPAG, bilaterally) were selected for analysis due to their relevance to vigorous escape and freezing responses, respectively ^16,17^. Three sections per animal were selected from the mid-rostrocaudal extent of the PAG, corresponding to bregma levels from -7.64 to -8.00 mm^24^, and images were acquired using a motorized confocal microscope (Zeiss Axio Imager Z1; Carl Zeiss AG, Germany). For each sample, z-stack series were acquired at 1 μm step intervals, with Iba1-labeled sections visualized at 40x objective magnification. During acquisition, the extended depth-of-focus (EDF) function was applied to generate two-dimensional projections from the z-stack images. All image datasets were coded to mask group identity before analysis.

Morphometric quantification was conducted using the Fiji distribution of ImageJ ^25^ combined with a dedicated macro (Microglial Morphology Analysis; https://github.com/BrainEnergyLab/Inflammation-Index)^26^, a robust high-throughput pipeline that enables efficient analysis of large numbers of cells while automating most processing steps unless otherwise specified by the user. To characterise microglial morphology, perimeter, number of branches and process architecture were evaluated on skeletonized cell masks using the Skeletonize and AnalyzeSkeleton plugins ^27^, while soma size was calculated using the FracLac plugin ^28^. In line with current recommendations for microglial morphometric analyses ^29^, all analysable microglial cells within the selected regions of interest were included in the quantification. To enable integrated comparisons of inflammation-associated morphological changes across distinct subregions and time points, these parameters were combined into a single inflammation index using an approach adapted from Clarke and collaborators ^26^. This method integrates the morphological features described above into a single composite metric by standardizing each parameter relative to the mean and standard deviation of the normoxia (control) group ^26^.

### 2.6 Experimental design

#### 2.6.1 Experiment 1 – Effects of the hypoxia challenge on animal behaviour and microglia morphology

The animals were pre-exposed to the experimental chamber one day prior to the experiment. During this session, each animal was placed individually in the chamber for 15 minutes while room air was injected at a flow rate of 4.5 L/min to allow adaptation to the environment and the sound of the air pump, aiming to mitigate neophobic reactions on the next day.

Twenty-four hours later, for the hypoxia challenge, each animal was placed in the chamber for an initial 5-minute period with room air flowing at a rate of 4.5 L/min for acclimatisation. Subsequently, N_2_ was flushed into the chamber at the same flow rate for 4 minutes to reduce the O_2_ concentration to 7%. After this period, the N_2_ infusion was interrupted, and the 7% O_2_ condition was maintained for an additional 6 minutes, during which we counted the number of jumps to the ceiling of the chamber, which was considered a panic index^30^. Test sessions were video-recorded and distance travelled during normoxia was subsequently analysed using AnyMaze software (Stoelting Co., Wood Dale, IL, USA). A separate group of animals remained under normoxic conditions (21% O₂) throughout the test.

At the end of the respiratory challenge, the animals returned to their respective home cages and after 1, 6, or 24h (n = 8/group) were anaesthetised and the brain of each rat was collected for the immunofluorescence analyses, as described above. Figure 1 (upper panel) shows an overview of the experimental design adopted in Experiment 1.

**Figure 1.**
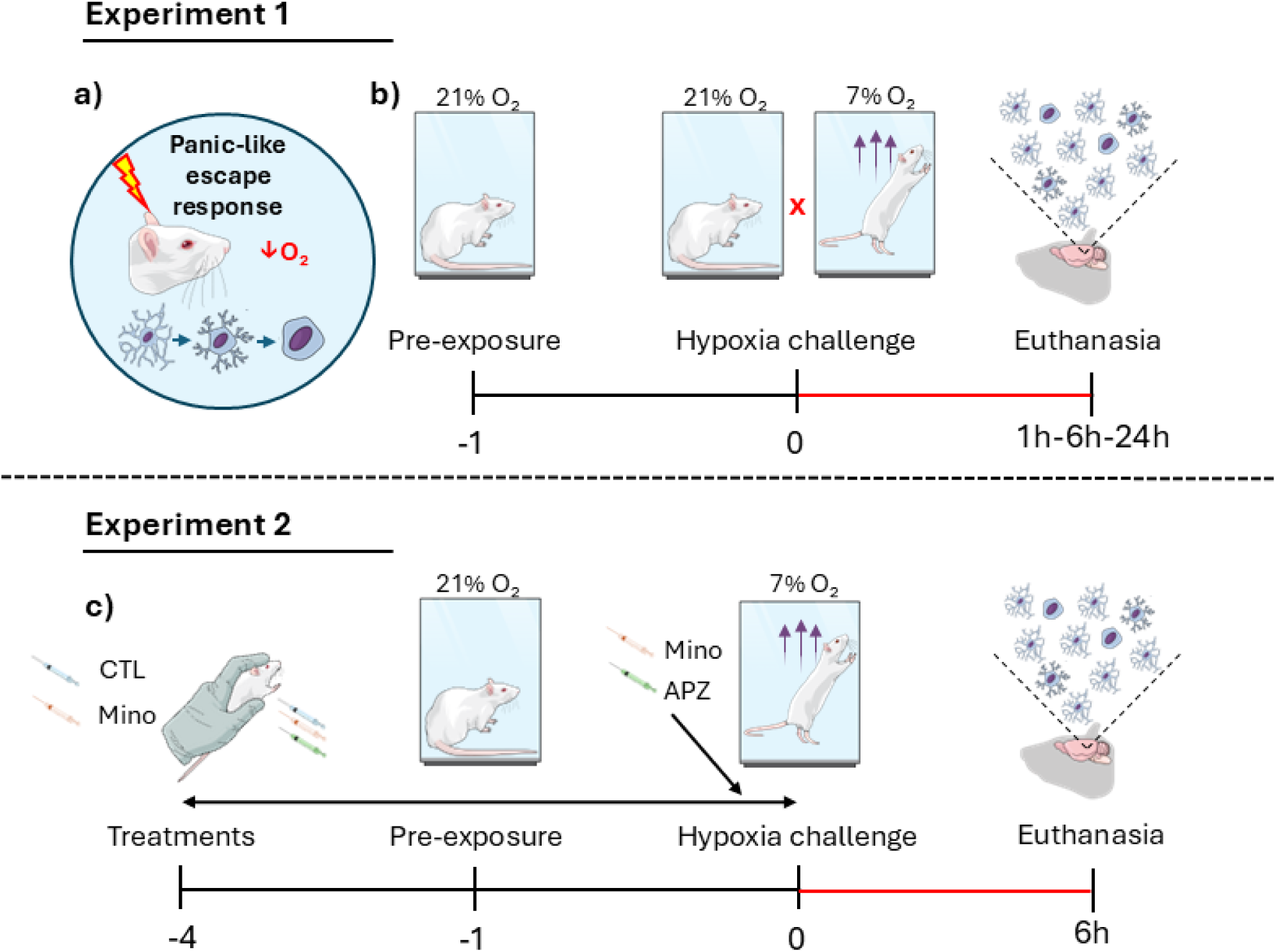
Schematic illustration of the experimental designs followed in Experiments 1 and 2. a) The hypoxia challenge model for investigating panic disorder (PD) was employed to assess time-dependent alterations in microglial morphology. b) Experiment 1: one day before testing, all animals were exposed to the gas chamber under standard conditions (21% O₂) for habituation. The following day, animals were subdivided into groups maintained either under normoxic conditions (21% O₂) or under hypoxic conditions (7% O₂). Each group (normoxia or hypoxia) was further divided and euthanised at 1, 6, or 24h after the challenge for microglial morphological analyses. c) Experiment 2: Animals were habituated to the gas chamber and exposed to hypoxia as described in experiment 1. In addition, they received an intraperitoneal (i.p.) injection of either vehicle (PBS; control group) or minocycline (Mino; 30 mg/kg ^31^) for five consecutive days. As a positive control, another group received an intraperitoneal injection of vehicle in the first four days and a single dose of alprazolam (Apz; 2 mg/kg ^33^) 30 minutes before testing on the fifth day. Animals were euthanised 6h after the hypoxia challenge for evaluation of microglial morphology.

#### 2.6.2 Experiment 2 – Effects of minocycline treatment on animal behaviour and microglia morphology

A new cohort of animals was randomly assigned to one of the following groups: minocycline (Mino; 30 mg/kg), alprazolam (Apz; 2 mg/kg) or vehicle. Minocycline or vehicle were administered intraperitoneally once daily for five consecutive days ^31,32^, whereas the alprazolam group received vehicle injections in the first four days and alprazolam on the test day ^33^. All treatments were administered in the morning (09:00-10:00 h), with the last injection given 30 min before the hypoxia challenge, which was performed as described in Experiment 1, except that no normoxia group was tested. Based on the microglial morphological changes observed in the previous experiment, the 6h post-hypoxia time point and the dmPAG were selected for microglial morphological analyses. Figure 1 (lower panel) shows an overview of the experimental design adopted in Experiment 2.

### 2.7 Statistical analysis

Data normality was assessed using the Shapiro-Wilk test. In Experiment 1, a two-way analysis of variance (ANOVA) was performed on the parameters: distance travelled, number of jumps, and microglia morphology with condition (normoxia or hypoxia) and time (1, 6 or 24h) as independent factors. In Experiment 2, a one-way ANOVA was applied to the same parameters, with treatment group (vehicle, Mino or Apz) as the independent factor. Post hoc comparisons were performed using Duncan’s test. Student’s t-test was used to compare microglial morphology between the dmPAG and vlPAG in Experiment 1, and between vehicle and Mino groups in Experiment 2. Data are expressed as mean ± standard error of the mean (SEM). Statistical analyses were carried out using SPSS (version 20.0), and figures were prepared with GraphPad Prism (version 9.0).

## 3 RESULTS

### 3.1 Hypoxia induces panic-like behaviour accompanied by time-dependent microglial morphological alterations

Figure 2 shows the behavioural effects observed in rats during the hypoxia challenge. Two-way ANOVA revealed significant effects of the respiratory condition [F(1,39) = 156.13; p < 0.001], with no significant effect of time [F(2,39) = 0.83; NS] nor an interaction between these two factors [F(2,39) = 0.83; NS] on the number of jumps. Notably, animals maintained under normoxic conditions did not exhibit this escape-like behaviour. In contrast, exposure to hypoxia elicited robust escape attempts, with no differences observed among hypoxia-exposed groups (Figure 2A). No change was observed in the animal’s locomotion (Figure 2B).

**Figure 2.**
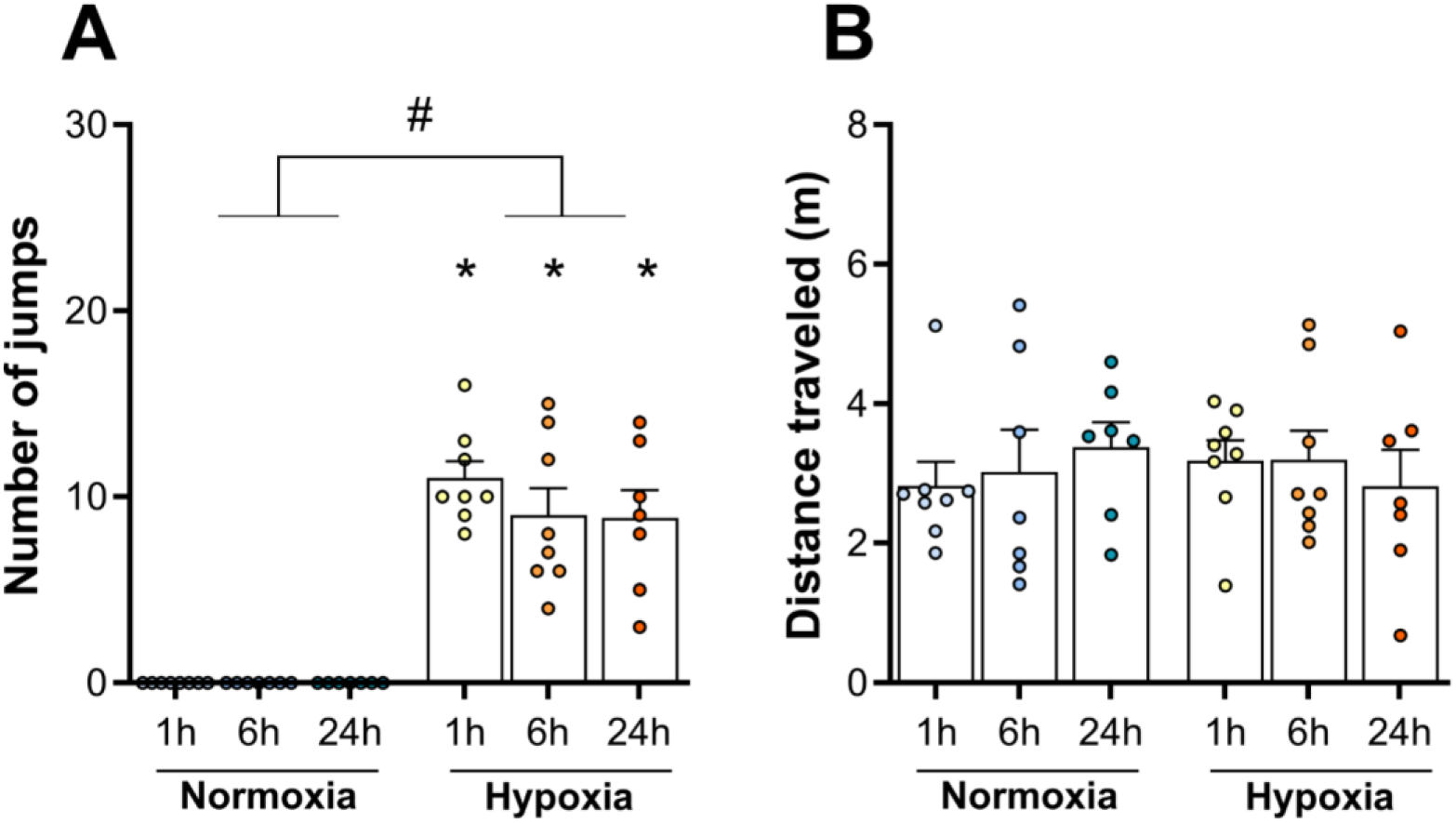
Effect (mean + S.E.M.) of normoxia (21% O₂) or hypoxia (7% O₂) exposure on: (A) number of jumps and (B) distance travelled. Data were analysed using two-way ANOVA. n = 7-8. ^#^*p* < 0.05 for condition effects; \**p* < 0.05 compared with the respective normoxia group.

Figure 3 shows representative images of microglial morphology in the dmPAG and vlPAG assessed at 1, 6, or 24h after the respiratory challenges.

**Figure 3.**
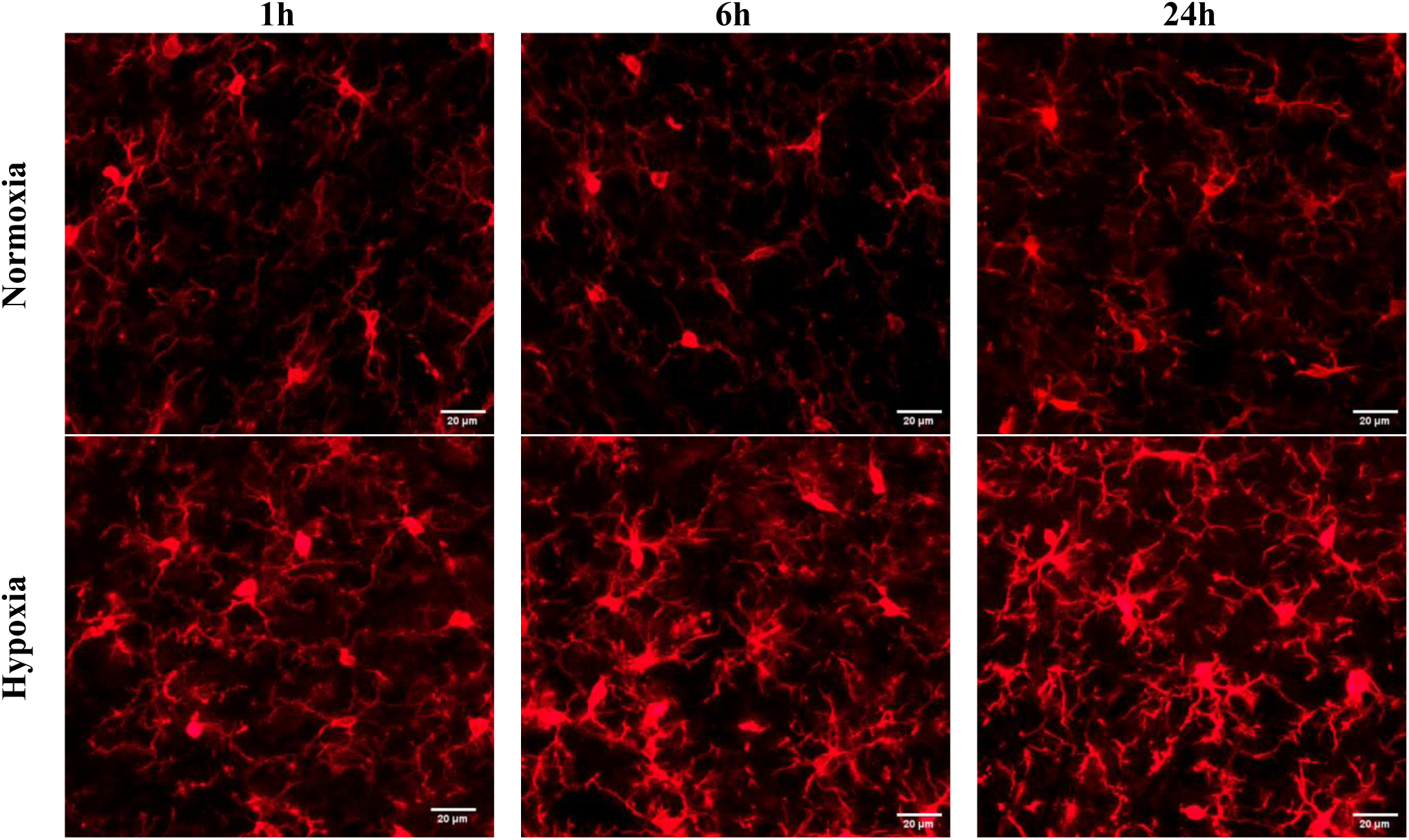
Representative fluorescence photomicrographs of microglia in the dorsomedial periaqueductal grey (dmPAG) obtained 1, 6, and 24 h after exposure to normoxic or hypoxic conditions. Microglia were labelled with anti-Iba1 and Alexa Fluor 594 antibodies and analysed by confocal microscopy to assess microglial morphology. 40x objective magnification. Scale bar: 20 μm.

In the dmPAG, two-way ANOVA revealed significant effects of condition [F(1,523) = 5.14; p < 0.05], time [F(2,523) = 6.85; p < 0.01], and their interaction [F(2,523) = 4.43; p < 0.05] on cell perimeter. Similar effects were observed on soma size, including condition [F(1,523) = 528.40; p < 0.001], time [F(2,523) = 59.14; p < 0.001], and their interaction [F(2,523) = 40.06; p < 0.001]. Exposure to hypoxia produced a time-dependent reduction in cell perimeter and soma enlargement (Figure 4A-B). A significant effect of the respiratory condition was also observed for the number of branches [F(1,523) = 46.75; p < 0.001], with hypoxic groups exhibiting decreased arborisation (Figure 4C). In the vlPAG, two-way ANOVA showed a significant effect of condition on cell perimeter [F(1,253) = 8.61; p < 0.01], characterised by reduced cell perimeters in hypoxia-exposed groups (Figure 4D). Significant effects on soma size were observed for condition [F(1,253) = 123.14; p < 0.001] and time [F(1,253) = 13.16; p < 0.01], showing progressive enlargement following hypoxic exposure (Figure 4E). In addition, a significant effect of condition was observed for the number of branches [F(1,253) = 20.92; p < 0.001], with reduced arborisation observed under hypoxic conditions (Figure 4F).

**Figure 4.**
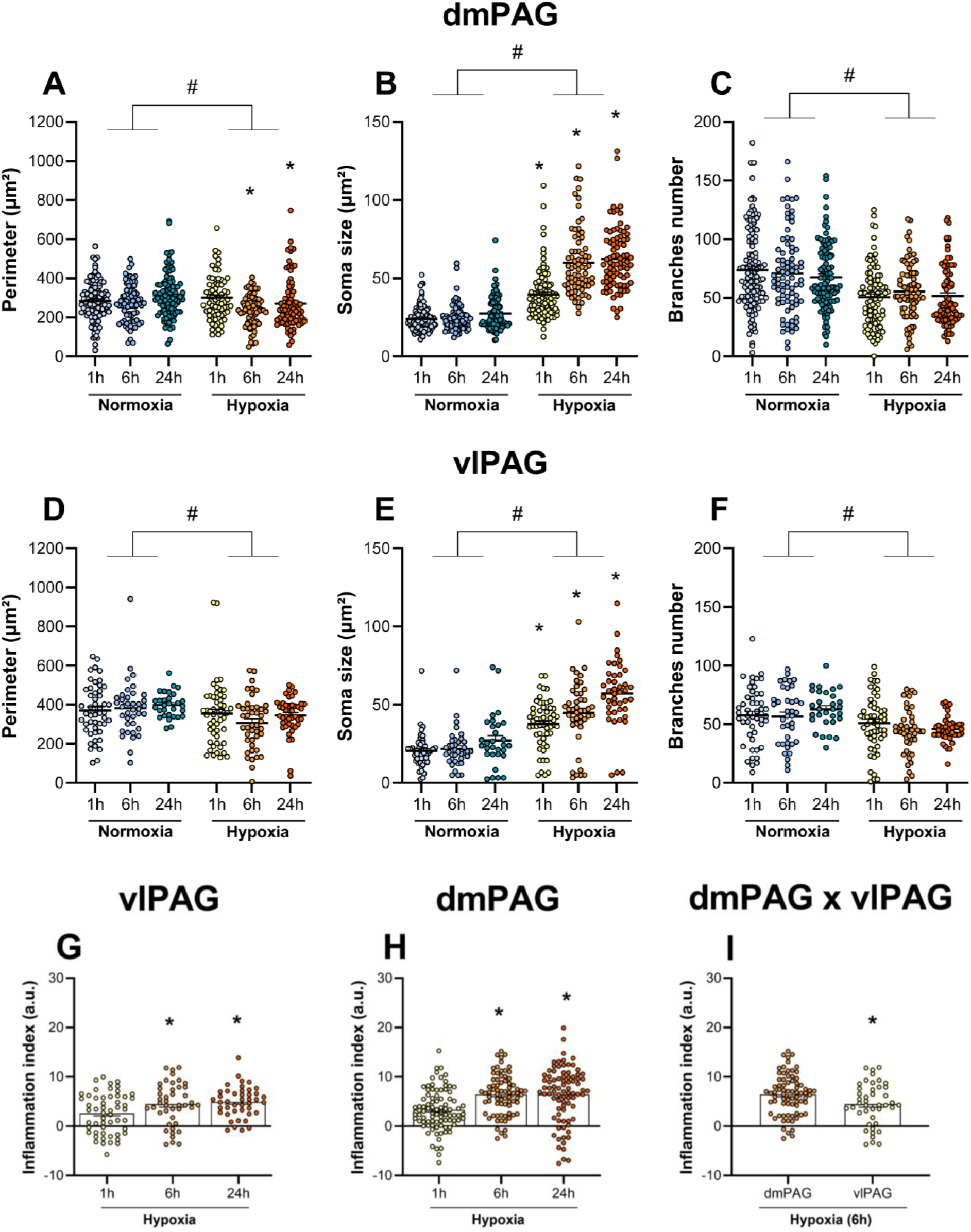
Effect (mean ± S.E.M.) of normoxia or hypoxia challenge on microglia morphology within the dorsomedial (dmPAG) and ventrolateral (vlPAG) periaqueductal grey columns. (A, D) Cell perimeter (B, E), soma size (C, F), number of branches (G, H, I), and inflammation index. Data were analysed using two-way ANOVA. A total of 69-121 (dmPAG) and 30-54 (vlPAG) cells were analysed from three midbrain sections per animal, with three animals included in each experimental group. Student’s t-test was used to compare differences between dmPAG and vlPAG; ^#^*p* < 0.05 for condition effects; \**p* < 0.05 compared with the respective normoxia group.

The inflammation index was calculated to enable comparisons across distinct PAG columns and time points. One-way ANOVA revealed that, following hypoxic exposure, both PAG columns investigated exhibited a subtle inflammatory response at 1h, which became more pronounced at 6h and remained elevated up to 24h [dmPAG: F(2, 223) = 8.92, p < 0.05; vlPAG: F(2, 137) = 5.03, *p* < 0.05], as shown in Figures 4G-H. At the 6h time point, a direct comparison between PAG columns revealed a higher inflammation index in the dmPAG [t = 2.45, df = 115, *p* < 0.01], as shown in Figure 4I.

### 3.2 Systemic minocycline treatment attenuates panic-like escape behaviour and microglial morphological alterations induced by hypoxia

Next, we investigated whether minocycline would inhibit escape expression during the hypoxia challenge and affect microglial morphology in the dmPAG. Alprazolam was chosen as a positive control for its known anti-panic effect ^33^. As shown in Figure 4, one-way ANOVA revealed that both sub-chronic (5 days) minocycline and acute alprazolam significantly reduced the total number of jumps exhibited during the hypoxia challenge [F(2,24) = 3.84, p < 0.05], without altering the distance travelled [F(2,24) = 0.46, NS] (Figure 5A-B).

**Figure 5.**
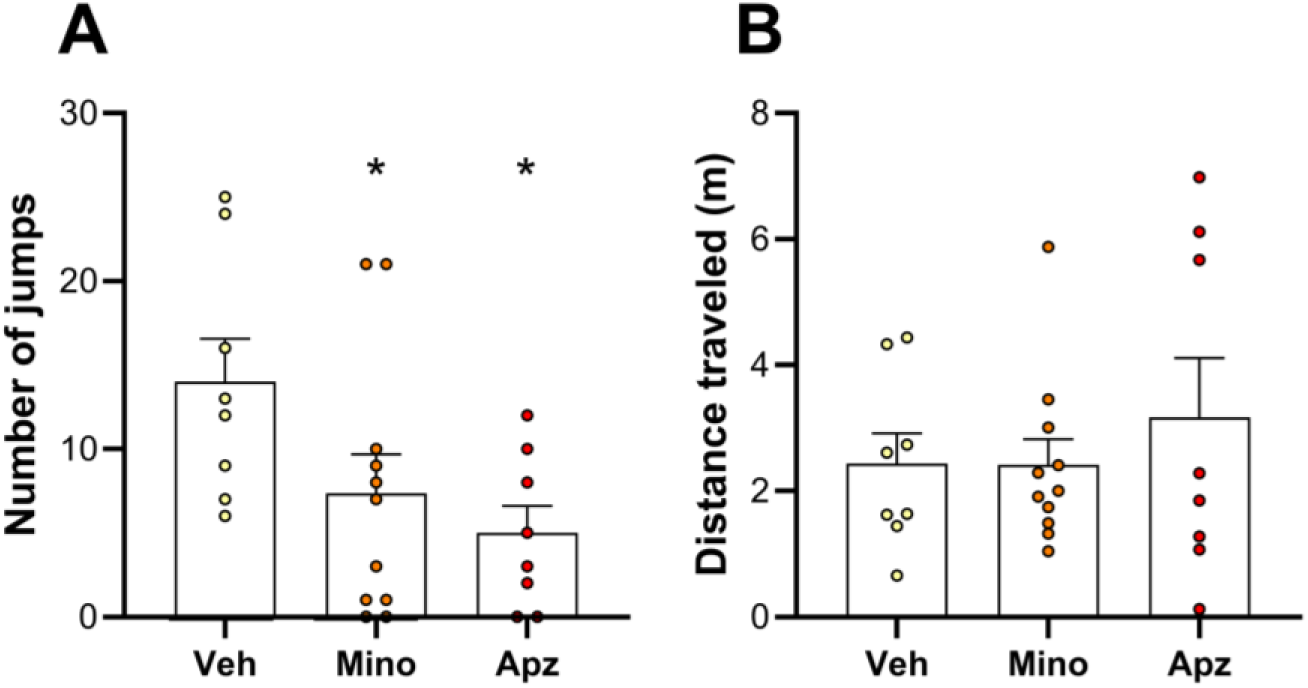
Effects (mean ± S.E.M.) of treatment with minocycline (Mino, 30 mg/kg, 5 days), alprazolam (Apz, 2 mg/kg, acute injection), or vehicle solution (Veh) in animals exposed to hypoxia on: (A) number of jumps, (B) distance travelled. Data were analysed using one-way ANOVA. n = 8-11. \**p* < 0.05 compared with the vehicle-injected group.

Figure 6A shows representative images of microglial morphology in the dmPAG 6 h after the respiratory challenge. Student’s t-test revealed significant effects of drug treatment on cell perimeter [t = 10.36, df = 161, p < 0.001], soma size [t = 6.13, df = 161, p < 0.001] and inflammation index [t = 8.75, df = 161, p < 0.001], but not on branch number [t = 1.64, df = 161, NS]. Minocycline administration prevented the hypoxia-induced increase in soma size and reduction in cell perimeter observed 6h after exposure. These effects were accompanied by a lower inflammation index compared with the vehicle-injected group, as shown in Figure 6B-E.

**Figure 6.**
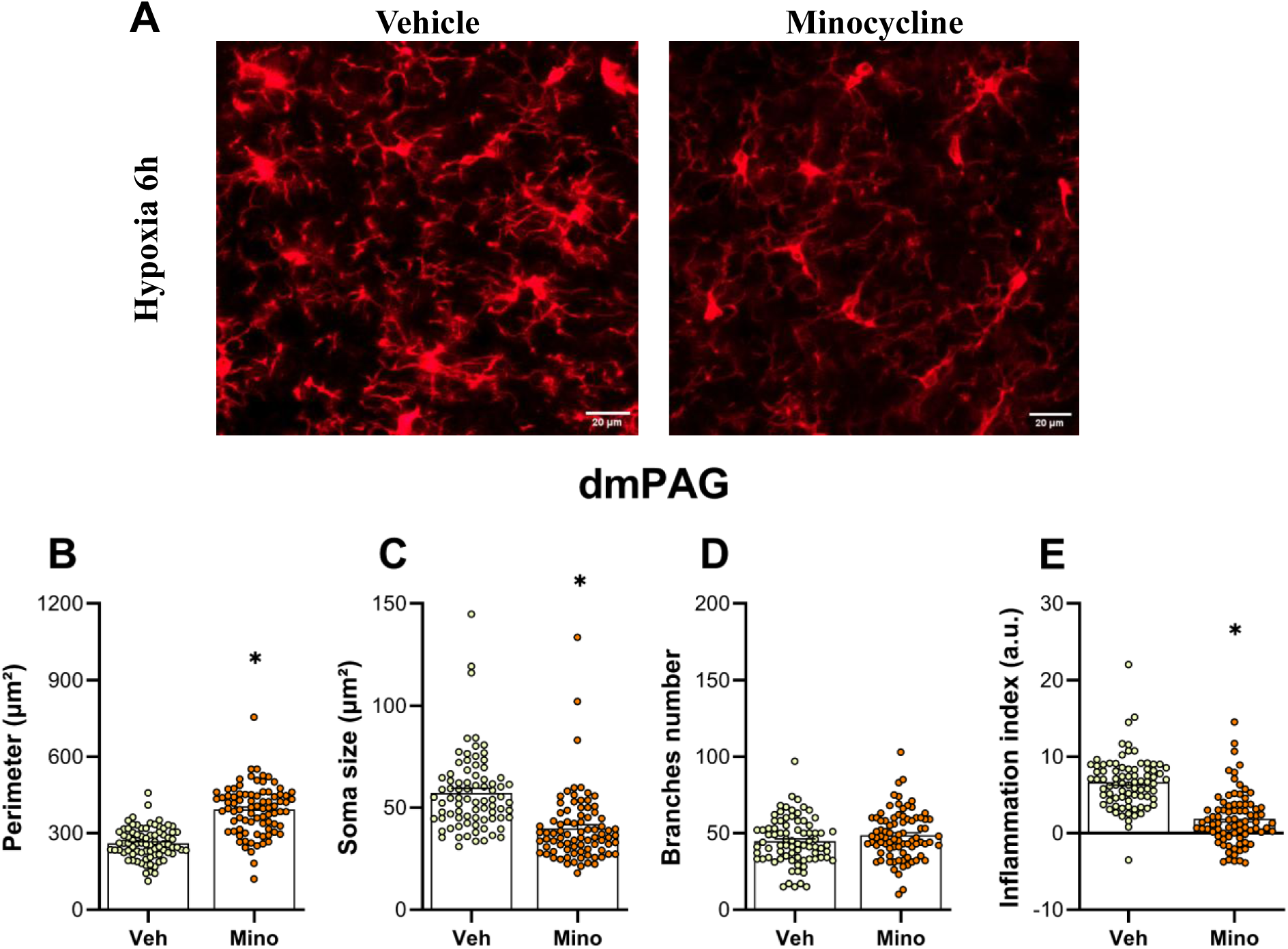
Effect (mean ± S.E.M.) of treatment with minocycline (30 mg/kg, i.p., 5 days) or vehicle solution on microglia morphology within the dorsomedial column of the periaqueductal grey (dmPAG), assessed 6 h after the hypoxic challenge. (A) Representative fluorescence photomicrographs of microglia (anti-Iba1 and Alexa Fluor 594) in the dorsomedial periaqueductal grey (dmPAG) 6h after exposure to hypoxic conditions in animals previously treated with vehicle solution (Veh) or minocycline, 40x objective magnification, scale bar 20 μm. (B) Cell perimeter, (C) soma size, (D) number of branches, and (E) inflammation index. Data were analysed using Student’s t-test. A total of 80-83 cells were analysed from three brain sections per animal, with three animals included in each experimental group. *p < 0.05 compared with the vehicle-injected group.

## 4 DISCUSSION

Despite advances in understanding PD pathophysiology, the broader cellular mechanisms underlying its susceptibility are not yet fully defined. Here, using hypoxia, a stimulus that evokes panic attacks in humans and panic-like behaviours in rats, we showed that microglia within a key PD-related area, the PAG, undergo a time-dependent shift towards an immunoresponsive phenotype. Notably, systemic administration of minocycline, similar to alprazolam, attenuated the expression of hypoxia-induced panic-like jumping behaviour, while preventing microglial activation in the dmPAG, highlighting a promising therapeutic avenue for PD.

The results of Experiment 1 showed that inhalation of low levels of O_2_ by rats consistently increases the number of escape attempts, independently of changes in overall locomotion, corroborating previous findings from our laboratory ^33–35^ and further supporting the reliability and reproducibility of this model. In a previous validation study, we also showed that treatment with first-line antipanic drugs used in clinical practice, including fluoxetine and alprazolam, inhibited this behavioural response, further supporting the translational value of this model ^33^.

A wealth of studies shows that both the dmPAG and the dorsolateral columns of the PAG (dlPAG) have been consistently implicated in the genesis/regulation of escape ^36–39^. For example, electrical or chemical stimulation of these PAG columns evokes vigorous escape reactions, similar to those observed under naturalistic conditions, such as confrontation with an approaching predator ^40^. Building on these findings, injecting interleukin-1 beta (IL-1β), a potent pro-inflammatory cytokine, into the cat dorsal PAG (dPAG) elicits defensive rage ^41^. This behaviour, which can also be evoked by dPAG electrical stimulation ^42^, reflects defensive reactions to actual or imminent threats, similar to escape responses. Based on this body of evidence, the next step in Experiment 1 was to investigate whether microglia, the resident immune cells of the central nervous system ^29^, were involved in panicogenic responses within this midbrain structure. We also investigated microglia involvement in the vlPAG, a column that has been associated with the regulation of freezing behaviour ^43^.

Microglia are specialised immune cells responsible for surveillance of the neural environment, playing essential protective and reparative roles ^29^. In their surveillant, healthy state, these cells display extensive surface area (perimeter) and highly arborised processes (branches), enabling continuous monitoring of the surrounding tissue ^29,44^. However, this morphological state can be altered by a range of factors, including tissue damage, infection, and reduced environmental oxygen concentrations, which promote extracellular acidosis. In response to this acidic environment, detected through acid-sensing receptors, growing evidence suggests that these cells can undergo morphological modulation ^14,45^. Consistent with this framework, the morphological changes observed in microglia in Experiment 1, especially within the dmPAG, which exhibited a higher inflammation index, are indicative of a shift towards an amoeboid phenotype, characterised by soma enlargement, reduced perimeter, and decreased branch number, all driven by tightly regulated cytoskeletal reorganisation. Importantly, this morphological transition is closely associated with an immunoresponsive state, typically marked by the release of pro-inflammatory cytokines and pro-oxidative mediators ^46^.

Furthermore, our findings align with prior rodent studies on hypoxia. For instance, Silva and collaborators ^31^ observed that adult male Wistar rats exposed to a similar hypoxic environment (8% O_2_) for a longer duration (3 hours) exhibited increased levels of pro-inflammatory cytokines, including IL-1β and TNF-α, within hypothalamic nuclei. In a protocol utilising 1-hour exposure to 8% O₂, Song and collaborators ^47^ reported similar cytokine elevations (e.g. TNF-α, IL-1β, and IL-6) in the cortex of male Sprague-Dawley rats, although behavioural outcomes were not assessed in either study. Consistent with these observations, the results from Experiment 1 suggest that microglia are recruited by short-term hypoxic exposure in the dmPAG and, to a lesser extent, the vlPAG, implicating these cells in the overall response to the respiratory challenge. Altogether, the present study identifies a novel association between immunoresponsive microglial states within these PAG columns and the behavioural consequences of exposure to panicogenic respiratory stimuli.

The literature addressing the kinetics of microglial morphology following injury or immunological challenges remains limited, with relatively few studies examining this process in detail. In one of the published studies, Norden and collaborators^48^ investigated microglial phenotypes in the hippocampus following a systemic immune challenge induced by lipopolysaccharide (LPS) administration in adult BALB/c mice. The authors reported that microglial morphological alterations, including enlarged somas, were detectable only 24 hours after endotoxin exposure and persisted for several days. By contrast, increases in pro-inflammatory cytokines, including IL-6, IL-1β, and TNF-α, occurred earlier, approximately 2 hours post-challenge. Despite employing a respiratory challenge rather than a standard pro-inflammatory agent such as LPS, this temporal dissociation closely aligns with the kinetics observed in our experiment. Specifically, we detected early microglial alterations 1 h after hypoxic exposure, whereas soma enlargement and significant reductions in cell perimeter and arborisation emerged predominantly within the dmPAG after 6 h and persisted for at least 24 h.

In Experiment 2, we administered minocycline intraperitoneally, a tetracycline-class antibiotic with recognised anti-inflammatory properties ^18–22^. Our findings demonstrate that minocycline produced a panicolytic effect, comparable to that observed following administration of alprazolam, a benzodiazepine previously shown by our group to effectively reduce escape attempts in this experimental paradigm ^33^. Previous studies have shown that minocycline can inhibit microglial activation across distinct brain regions, including the hypothalamus and hippocampus, often accompanied by reductions in anxiety-like behaviour and improvements in learning and memory performance ^20,31,49^. Regarding morphological aspects, pre-treatment with minocycline (25 mg/kg for 4 days) before LPS injections has been shown to suppress LPS-induced microglial transition towards an amoeboid phenotype in Sprague Dawley rats ^50^. Interestingly, a similar dose, route, and duration of minocycline administration (i.p., 30 mg/kg, 5 days) to that used in the present study were previously shown to reduce hypothalamic levels of IL-1β and TNF-α five days after hypoxia challenge (8% O₂ for 3h) in Wistar rats ^31^.

Although several mechanisms have been proposed to explain the effects of minocycline on microglial phenotype^51^, one pathway has attracted particular attention. Specifically, modulation of the Janus kinase/signal transducer and activator of transcription (JAK/STAT) signalling pathway appears to be one of the most relevant mechanisms. Minocycline has been shown to suppress STAT1-associated pro-inflammatory responses while favouring a more reparative phenotype potentially mediated by STAT6 activation. This shift is accompanied by reduced production of pro-inflammatory cytokines, together with enhanced expression of anti-inflammatory mediators ^51^. Nevertheless, further studies are required to confirm the involvement of these mechanisms in the present model.

More specifically in relation to PD, our group has recently demonstrated the therapeutic potential of minocycline in experimental models of respiratory panic, as well as in human patients diagnosed with this disorder ^9^. Adult male C57BL/6 mice exposed to hypercapnia (20% CO₂) – a condition that similarly evokes a robust jump escape response only in mice^8^ – were assessed for panic-like behaviours and microglial morphology in the locus coeruleus after subchronic minocycline treatment (40 mg/kg i.p., 14 days). The findings revealed reduced microglial arborisation 6h following hypercapnic exposure, alongside a panicolytic effect of minocycline, although the direct impact of treatment on microglial morphology was not evaluated. Notably, in humans diagnosed with PD, treatment with minocycline, compared with the benzodiazepine reference drug clonazepam ^52^, reduced the severity of CO₂-induced panic attacks while modulating immune responses, as evidenced by decreased IL-2sRα and increased IL-10 levels ^9^. Building on this evidence, the present study advances the field by demonstrating that minocycline effectively reduces the microglial transition towards an amoeboid phenotype after 6h of hypoxia, highlighting its potential not only to attenuate panic-like behaviours but also to modulate inflammatory processes associated with an immunoresponsive phenotype within midbrain regions implicated in PD pathophysiology. Future studies incorporating both sexes and complementary molecular markers of microglial activation will be important to further validate and extend these observations.

## 5 CONCLUSION

This study demonstrates that hypoxic challenge induces time-dependent morphological changes in microglia, which are significantly more pronounced in the dmPAG, a midbrain structure critical for generating panic-like responses and regulating defensive behaviour, than in the vlPAG. Our findings indicate that hypoxia shifts microglia towards an immunoresponsive phenotype. These alterations are evident as early as 6 hours after hypoxic exposure and persist for at least 24 hours. Furthermore, minocycline administration attenuates both the behavioural and microglial responses to hypoxia, highlighting microglial modulation as a promising therapeutic strategy for PD.

## ACKNOWLEDGMENTS

The financial support was provided by FAPESP (2020/15050-7, <u>2021/08968-0</u>, <u>2024/13469-1</u> to AR-G, 2025/28895-9 to NCC, and 2017/19731-6 to SFL); by the CNPq (140323/2022-8 to JMNS); by the AMS Springboard award SBF005\1102 and the MRC Career Development Award MR/T031115/1 to VM. SFL and HZJ are recipients of CNPq Research Productivity Fellowship. We also thank Afonso Paulo Padovan for technical support.

## Ethics approval statement

It was approved by the Ethics Committee on the Use of Animals under protocol (1134/2022).

